# Development of a Xylose-Inducible Promoter and Riboswitch Combination System for Manipulating Gene Expression in *Fusobacterium nucleatum*

**DOI:** 10.1101/2023.04.24.538132

**Authors:** GC Bibek, Peng Zhou, Arindam Naha, Jianhua Gu, Chenggang Wu

**Affiliations:** Department of Microbiology & Molecular Genetics, the University of Texas Health Science Center, Houston, TX, USA; Houston Methodist Hospital Research Institute, Houston, TX

**Keywords:** *Fusobacterium nucleatum*, a xylose-inducible system, riboswitch, essential genes, The type I signal peptidase

## Abstract

Inducible gene expression systems are important for studying bacterial gene function, yet most exhibit leakage. In this study, we engineered a leakage-free hybrid system for precise gene expression controls in *Fusobacterium nucleatum* by integrating the xylose-inducible expression system with the theophylline-responsive riboswitch. This innovative method enables concurrent control of target gene expression at both transcription and translation initiation levels. Using luciferase and the indole-producing enzyme tryptophanase (TnaA) as reporters, we demonstrated that the hybrid system displays virtually no observable signal in the absence of inducers. We employed this system to express FtsX, a protein related to fusobacterial cytokinesis, in an *ftsX* mutant strain, unveiling a dose-dependent manner in FtsX production. Without inducers, cells form long filaments, while increasing FtsX levels by increasing inducers concentrations led to a gradual reduction in cell length until normal morphology was restored. Crucially, this system facilitated essential gene investigation, identifying the signal peptidase *lepB* gene as vital for *F. nucleatum*. LepB’s essentiality stems from depletion, affecting outer membrane biogenesis and cell division. This novel hybrid system holds the potential for advancing research on essential genes and accurate gene regulation in *F. nucleatum*.

## IMPORTANCE

*Fusobacterium nucleatum*, an anaerobic bacterium prevalent in the human oral cavity, is strongly linked to periodontitis and can colonize areas beyond the oral cavity, such as the placenta and gastrointestinal tract, causing adverse pregnancy outcomes and promoting colorectal cancer growth. Given *F. nucleatum*’s clinical significance, research is underway to develop targeted therapies to inhibit its growth or eradicate the bacterium specifically. Essential genes, crucial for bacterial survival, growth, and reproduction, are promising drug targets. A leak-free inducible gene expression system is needed for studying these genes, enabling conditional gene knockouts and elucidating the importance of those essential genes. Our study identified *lepB* as the essential gene by first generating a conditional gene mutation in *F. nucleatum*. Combining a xylose-inducible system with a riboswitch facilitated the analysis of essential genes in *F. nucleatum*, paving the way for potential drug development targeting this bacterium for various clinical applications.

## INTRODUCTION

The human oral cavity is home to hundreds of bacterial species (1–3) that predominantly exist within polymicrobial biofilm rather than in a planktonic state (4, 5). Two main types of biofilms can be found in the mouth: the supragingival biofilm and the subgingival biofilm. Supragingival biofilm is found at or above the gingival margin. In contrast, subgingival biofilms are located in the so-called periodontal pocket, where they come in close contact with gingival tissue. Their presence has a strong correlation with periodontal disease (6). Approximately 400 bacterial species inhabit the subgingival biofilm, physically and metabolically interconnected to form a dynamic, well-organized multispecies microbial community commonly known as dental plaque (7, 8).

One bacterial species in dental plaque is *Fusobacterium nucleatum*, a Gram-negative, spindle-shaped anaerobe (6). *F. nucleatum* plays a crucial role in the development and maturation of subgingival biofilm (9), as it has the ability to aggregate with other oral bacteria, including early colonizers such as *Actinomyces* spp. and oral streptococci and late colonizers such as *Porphyromonas gingivalis*, *Tannerella forsythia*, and *Treponema denticola* (8). Researchers believe that *F. nucleatum* acts as a bridge between these two groups since early colonizers usually do not interact physically with late colonizers (8, 10). In vivo experiments have demonstrated that *F. nucleatum* is located in the middle layer of dental plaque, physically connecting early and late colonizers (11). This physical connection enhances signal communication and metabolic exchange between plaque bacteria, making *F. nucleatum* critical not only for the structural organization of dental plaque but also for the metabolic network (4, 12, 13).

*F. nucleatum*’s ability to bridge connection is attributed to two primary adhesins on its cell surface: RadD (14) and Fap2 (15). Fap2 plays a vital role in the coaggregation of fusobacteria with certain strains of *P. gingivalis* (15) and *Enterococcus faecalis* (16). In contrast, RadD exhibits a broader range of interactions, enabling *F. nucleatum* to aggregate with various Gram-positive early colonizers, such as *Actinomyces oris*, *Streptococcus oralis*, *Streptococcus cristatus*, and *Streptococcus gordonii* (14, 17, 18). Furthermore, RadD assists in fusobacterial interactions with *Streptococcus mutans* (19), *Clostridioides difficile* (20), *Staphylococcus aureus* (21), *Aggregatibacter actinomycetemcomitans* (17) and oral fungus *Candida albicans* (22). Our lab’s research focuses on *radD* gene regulation and biogenesis.

RadD, an autotransporter protein, is anticipated to be secreted via the type Va secretion pathway(14). This multi-domain protein consists of an N-terminal signal sequence, a C-terminal β-domain, and a secreted passenger domain that carries the protein’s function in between (23). The signal sequence supposedly interacts with the Sec or Tat translocon to facilitate transport across the inner membrane, after which the signal peptidase removes it (24). In *E. coli*, LepB is a type 1 signal peptidase responsible for extracting the signal peptide of protein precursors, allowing proteins to mature and fold properly as they are translocated across the bacterial cytoplasmic membrane (25, 26).

The *F. nucleatum* strain ATCC 23726 genome contains a gene locus annotated as *lepB* (27) (HMPREEF0397_0712, accessible at http://img.jgi.doe.gov/). We reasoned that this LepB might interact with RadD’s signal peptide. Should *lepB* be deleted, the presentation of RadD on the cell surface could be diminished or lost, compromising *F. nucleatum*’s ability to aggregate with its partners. To investigate this, we attempted to create *lepB* in-frame deletion mutants using a *galk*-based method recently established in our laboratory (28), but it proved unsuccessful, suggesting that the deletion of *lepB* was lethal to the cell. This led us to hypothesize that the *lepB* gene is essential in *F. nucleatum*. An essential gene can be deleted from the chromosome if a second wild-type copy of this gene is actively expressed within the cell. And controlling the expression of this second target gene copy typically involves using a controlled gene inducible system (29–31). Ideally, these inducible systems should be leak-free. The rationale behind this is that certain gene products can satisfy the growth needs of their host, even at very low concentrations, owing to their high activity levels; a leak-free inducible system can result in the total depletion of the target gene product, thereby aiding in comprehending the target gene’s importance.

In *F. nucleatum*, two gene-inducible systems are currently in use: the tetracycline-controlled expression system (27) and the theophylline-sensed riboswitch (32). While the tetracycline-controlled system is effective, it has a drawback, as the inducers it employs (tetracycline or anhydrotetracycline) are toxic to some *F. nucleatum* strains. Furthermore, this system exhibits some leakage, which was discovered when we applied it to another oral bacterium, *A. oris* (29). Not only does the tetracycline system leak (33), but the theophylline-responsive riboswitch also has a slight leak (34). To eliminate these leaks, we combined the two systems to create a leak-free system to study the essentiality of housekeeping sortase *srtA* gene in *A. oris* (29).

Inspired by this approach, in this study, we developed a novel leak-free expression system that combines a *Clostridioides difficile*-derived xylose-inducible system (35) with theophylline-responsive riboswitch. To test this system, we used it to control the expression of the FtsX protein, which plays a role in cell division. The lack of the *ftsX* gene results in elongated filamentous cells (28). By increasing concentrations of the inducer(s) (xylose and/or theophylline), we observed a dose-dependent rise in FtsX levels, leading to a progressive reduction in cell length until it matched with the wild-type cell length. Notably, without inducers, the cell phenotypes were identical to the *ftsX* mutant strain, illustrating tight control over target gene expression. Subsequently, we employed this system to control the ectopic expression of the *lepB* gene, successfully generating a conditional *lepB* mutant strain. In the absence of inducers, cells perished due to incorrect outer membrane biogenesis and disrupted cell division by *lepB* depletion. This is the first instance of constructing conditional mutant strains for studying an essential gene in *F. nucleatum*.

## RESULTS

### Construction of a xylose-inducible and riboswitch-coregulated expression system for *F. nucleatum*

In this study, we aimed to create an inducible system for precise gene expression control and investigation of essential genes in *F. nucleatum*. We had previously developed a leak-free hybrid gene expression system that combined the tetracyclines-inducible (Tet) system and theophylline-sensed riboswitch for use in the oral bacterium *A. oris* (29). Our initial plan was to adapt this system for *F. nucleatum*; However, we found that the *F. nucleatum* strain we were working with was susceptible to the inducer anhydrotetracycline (ATc). As a result, we opted for an alternative inducible system known as the xylose-inducible system for two reasons: (1) *F. nucleatum* does not metabolize xylose, and (2) the inducer xylose does not affect *F. nucleatum* growth at concentrations below 1%. Significant growth inhibition occurs only when the xylose concentration exceeds 3% (Fig.S1). The xylose-inducible system (XIS) has been successfully employed for various Gram-negative bacteria, including *Pseudomonas fluorescens* (36) and *Caulobacter crescentus* (37), as well as Gram-positive species such as *Bacillus subtilis* (38), *C. difficile* (35), *Lactococcus lactis* (39), *Staphylococcus xylosus* (40), and *Streptomyces lividans* (41). This system primarily consists of *xylR*, the intergenic *xyl* operator sequence *xylO*, and the divergent *xyl_BA_* promoter. In the absence of the inducer xylose, XylR binds to *xylO*, blocking the transcriptional initiation of the target gene. In contrast, when xylose is present, it directly binds to XylR, causing XylR to detach from *xylO* and initiating target gene expression (Fig. 1A and 1B). We chose to work on the xylose-inducible system coming from *C. difficile* because *F. nucleatum* is phylogenetically closer to the *Clostridioides* genus (42), and both are low GC content, despite *F. nucleatum* being a Gram-negative bacterium and *C. difficile* a Gram-positive one. Using the *Clostridioides* -derived xylose-inducible system addresses potential codon bias issues.

**Figure 1:**
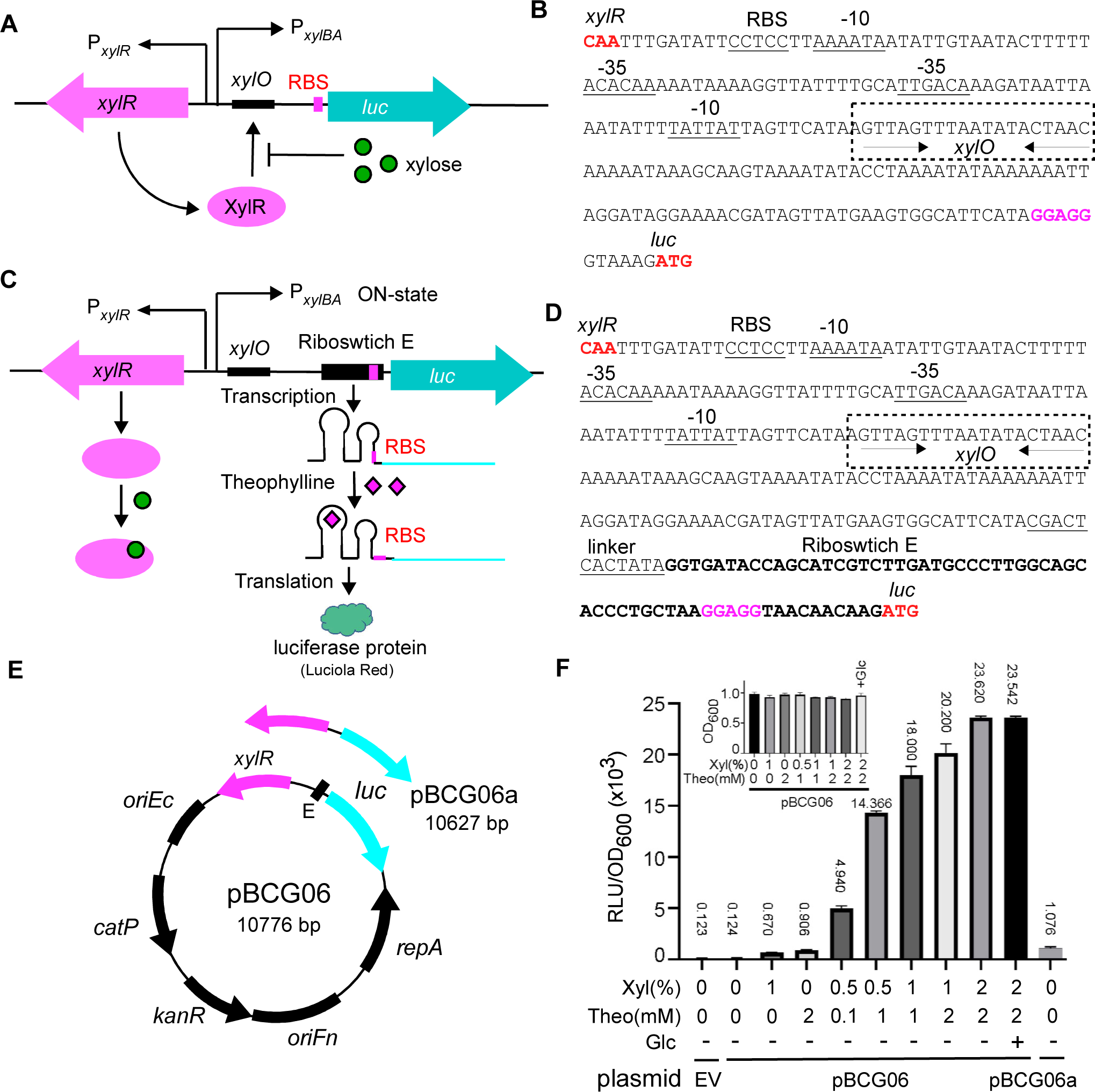
Construction and characterization of xylose and riboswitch-based dual inducible system in *F. nucleatum*. **(A)** Schematic diagram of assembling and working mode of the xylose-inducible system developed in this study to regulate reporter gene expression. This system was constructed by combining xylose repressor gene *xylR*, its operator sequence *xylO*, and the strong constitutive promoter P*_xylAB_*, the divergent promoter to drive the expression of Luciola Red luciferase (*luc*) gene reporter. The XylR is expressed constitutively from P*_xylR_*. It binds to the XylR operator (schematically depicted as a black) and blocks *luc* expression from the P*_xylAB_* promoter in the absence of xylose. XylR is released from the operator with xylose, initiating the *luc* expression. **(B)** The intergenic promoter region of the divergently transcribed *xylR* and *luc* in this xylose-inducible system. The translation initiation codon and the ribosomal bind site (RBS) are in bold red. The -10 and -35 sequences are shown as underlined sequences. The *xylO* element is highlighted in the box with dashed lines. Palindromic sequences in *xylO* are indicated by arrows. **(C)** Schematic diagram of assembling and working mode of the xylose and riboswitch-based dual inducible system developed in this study. The dual control system contains *xylR*, *xylO*, P*_xylAB,_* and a theophylline-responsive synthetical riboswitch E unit. The riboswitch is inserted to replace the RBS in the xylose-inducible system **(A)**. The reporter *luc* gene expression is initially repressed by XylR and is only activated in the presence of xylose. In the latter case, the *luc* gene is transcribed, but the mRNA is not translated in the absence of the riboswitch signal theophylline (RBS is hidden in a step-loop structure in riboswitch). In the presence of theophylline, the riboswitch changes its conformation and exposes the RBS so the ribosome can bind, resulting in synthesizing the luciferase protein. **(D)** The intergenic promoter region of the divergently transcribed *xylR* and *luc* in this dual inducible system. The linker region (underlined) and riboswitch E sequence were positioned as indicated in Figure (C). **(E)** Genetic maps of the xylose and riboswitch-based dual inducible expression reporter plasmid pBCG06 and the xylose-inducible alone system expression plasmid pBCG06a. Both plasmids are derivatives of *E. coli-F. nucleatum* shuttle vector pCWU6. Feature depicted: The *xylR-PxylAB* is from the vector pXCH, a xylose-inducible gene expression vector for *Clostridium perfringens* (35); *luc* encodes a Luciola Red luciferase protein that is codon optimized for expression in low-GC bacteria. *oriFn* and *repA*, the replication region; *oriEc*, replication region of the *E. coli* plasmid pBR322; *catP*, the chloramphenicol acetyltransferase gene, conferring resistance to thiamphenicol in *F. nucleatum* or chloramphenicol in *E. coli*; *KanR* is kanamycin resistance cassette from pJRD215. **(F)** Tunable induction of luciferase protein from P*_xylAB_* and the riboswitch control with different inducer(s) concentrations. The overnight cultures of *F. nucleatum* strain ATCC 23726 carrying pCWU6 (empty vector, EV), pBCG06, or pBCG06a were diluted 1:20 into TSPC. When cultures grew to OD600 ∼0.65, the inducers were added with various concentrations, as indicated. A tested culture received 20 mM glucose to assess its impact on the inducible system. After 2 hours of further incubation, 100 µl aliquots of each culture were taken out for luciferase assay. Luciferase activity (RLU) was normalized with cell density (OD_600_). Each culture’s mean RLU/OD_600_ was indicated above the corresponding column. The inserted graph showed the OD_600_ value of the cultures when the luciferase assay was performed. Data represent the means of standard deviations of the results of triplicate biological experiments.

All xylose-inducible systems that have been developed display varying degrees of leakiness, and the system derived from *C. difficile* is not an exception to this trend. To improve this system, we integrated a theophylline-responsive riboswitch E element (abbreviated as E, see Figure 1E), previously used for gene expression control in *F. nucleatum* (29), into the region between the target gene and the xylose-inducible promoter. Essentially, this riboswitch replaced the original ribosome-binding site (RBS) on the *xyl* promoter (*P_xylBA_*) (Figures 1C and 1D). The theophylline riboswitch introduces an additional layer of gene expression regulation at the translational level, distinct from the xylose-inducible system, which modulates gene expression at transcription initiation and requires the accessory protein (XylR) for induction. Suppose the xylose-inducible system (XIS) produces leaky mRNA in the absence of the inducer xylose. In that case, the translation will not occur without theophylline since the mRNA’s ribosome-binding site (RBS) is hidden within a step-loop structure of the riboswitch. A conformational change that exposes the RBS and initiates translation occurs only when theophylline binds to its aptamer domain in the riboswitch (Figures 1C and 1D). With this dual transcriptional (XIS) and translational (riboswitch E) control system in place, we expect to observe no or negligible leak expression when using it for gene expression in *F. nucleatum*.

To evaluate this expectation, we created the plasmid pBCG06(*xylR-E-luc*), which was specifically designed to express the Luciola red luciferase gene (*luc*) under the control of the XIS-riboswitch (Figure 1E). The Luciola red luciferase offers greater sensitivity and brighter luminescence than Firefly luciferase (43), which we previously used as a reporter for gene expression monitoring in *F. nucleatum* (17). We introduced the plasmid into *F. nucleatum* and cultured the cells. When the cultures’ OD_600_ values reached 0.65, we added various concentrations of inducers (xylose or theophylline alone or in different combinations). We allowed the cells to grow for two hours in an anaerobic chamber. After two hours, we removed the fusobacterial cultures from the anaerobic chamber, collected three 100 µl samples from each tested culture, and added them to a 96-well plate on the lab bench. We mixed each sample with 25 µl of 1 mM D-luciferin suspended in 100 mM Citrate buffer (pH 6.0) and vigorously pipetted 15 times aerobically with a multi-channel pipette. This process ensured that oxygen, which is required for luciferase activity, was available within the cells. Subsequently, we measured luciferase activity using a Microplate Luminometer. Fusobacterial cells with pCWU6, the empty vector (EV), were used as a negative control. In the absence of inducers, we noted minimal luciferase activity, comparable to the control culture grown with the fusobacterial cells containing an empty plasmid (EV) (column 1 vs. column 2 in Figure 1F). This indicated that the signal detected without inducers represented background noise, suggesting no leak in this system. Upon adding xylose alone, we observed a slight increase in luciferase activity, implying a minor leakiness in the riboswitch (Figure 1F, column 3). Likewise, when only theophylline was added, we detected an increased signal. This demonstrated that the xylose-inducible system also exhibited leakiness, with its degree of leakiness somewhat higher than that of the riboswitch (Figure 1F, column 4). This result is in line with the outcomes from another parallel control experiment we carried out. In that experiment, we generated a pBCG06a plasmid in which the expression of the luciferase gene was controlled exclusively by the xylose-inducible system (Figures 1A and 1E). We cultivated fusobacterial cells with this plasmid under the same conditions applied to the cells with pBCG06- and evaluated luciferase activity without the presence of the xylose inducer. The luciferase activity was at a particular level (Figure 1F, the last column), similar to cells containing the pBCG06 plasmid when only theophylline was used. This evidence strongly reinforces the notion that the xylose-inducible system inherently exhibits leakiness. However, when the riboswitch was integrated with the xylose-inducible system, it effectively mitigated this leakiness. As the concentrations of both inducers, xylose, and theophylline, progressively increased, the luciferase activity exhibited a positively correlated rise.

*F. nucleatum* has been reported to utilize glucose and possess a CcpA protein (43), often resulting in catabolite repression of sugar utilization in certain bacteria. To examine if glucose affects this hybrid system, we added 20 mM glucose along with 2% xylose and 2 mM theophylline to a culture. The luciferase activity in cells grown with glucose was almost identical to that in cells without glucose, indicating that this system remained unaffected by glucose (compared to Figure 1F, the last two columns). It is important to note that the xylose and theophylline used in these experiments did not alter bacterial growth, as shown in Figure 1F (the inserted graph).

*F. nucleatum* is a heterogenous species consisting of four subspecies: subsp. *nucleatum*, subsp. *vincentii*, subsp. *polymorphum* and subsp. *animalis* (44). The distribution of these subspecies within the human body varies. For instance, *F. nucleatum* subsp. *nucleatum* (FNN) is frequently isolated from cases of periodontal disease (45), while subsp. *animalis* (FNA) is most commonly associated with colorectal cancer tissue (46). Our current research focuses on the FNN strains. But it is intriguing to know if the XIS-riboswitch system exhibits comparable expression effects on other subspecies strains. To investigate this, we introduced pBCG06 into FNA strain ATCC 51191. The results indicated that except for a modest decrease in luciferase activity compared to FNN, FNA containing pBCG06 displayed a similar dose response as observed in FNN. Importantly, no leak expression was detected (Figure S2). These results strongly suggest that the XIS-riboswitch hybrid system can offer a leak-free advantage in *F. nucleatum*, responding to inducers in a dose-dependent manner.

### Utilize indole as a reporter to further validate the use of this XIS-riboswitch system

To assess the generality of the XIS-riboswitch system response across different target genes, we generated another plasmid construct, pBCG07. This construct enables the expression of the tryptophanase gene (*tnaA*) derived from FNN, governed by the XIS-riboswitch (Figure 2A). The *tnaA* gene has not been used as a reporter before. TnaA is responsible for indole production (32). When Indole reacts with para-Dimethylaminobenzaldehyde (DMAB) under acidic conditions, it forms the red dye rosindole. The dye is easily observable, without needing a complex machine, unlike fluorescence protein GFP and mCherry, and its quantity is proportional to indole levels. As rosindole concentration increases, the red color’s intensity changes, enabling indole estimation. Additionally, the indole assay provides a quantitative method for detecting indole levels (47). Thus, Indole could serve as a gene reporter in *F. nucleatum* and help evaluate our XIS-riboswitch inducible construct. To explore this possibility, we transformed pBCG07 into a recently generated *tnaA* mutant strain of FNN (32). The transformant *F. nucleatum*/pBCG07 was cultured for 15 h (cultures reached the late logarithmic phase) in a TSPC medium containing 0 or various concentrations of xylose and theophylline combinations. Next, 500 µl of Kovacs’s reagent was added to each culture tube without shaking. As shown in Figure 2B, indole production in these cultures was signaled by a yellow, pink, to red color shift. In the absence of inducers, indole production was undetectable (yellow); however, indole production became evident when xylose and theophylline were added individually or combined. Introducing 1% xylose yielded a light color, while 2 mM theophylline yielded a red hue. As the concentration of both inducers increased, so did the intensity of the red color. This escalation in red color intensity, reflecting elevated indole levels, was associated with the rising total concentrations of the two inducers. The quantification of indole levels in the culture (Figure 2C) further confirmed this observation. These findings align with those acquired using luciferase as a reporter, signifying that the XIS-riboswitch system operates without leakage in *F. nucleatum* and triggers gene expression in a dose-responsive manner.

**Figure 2:**
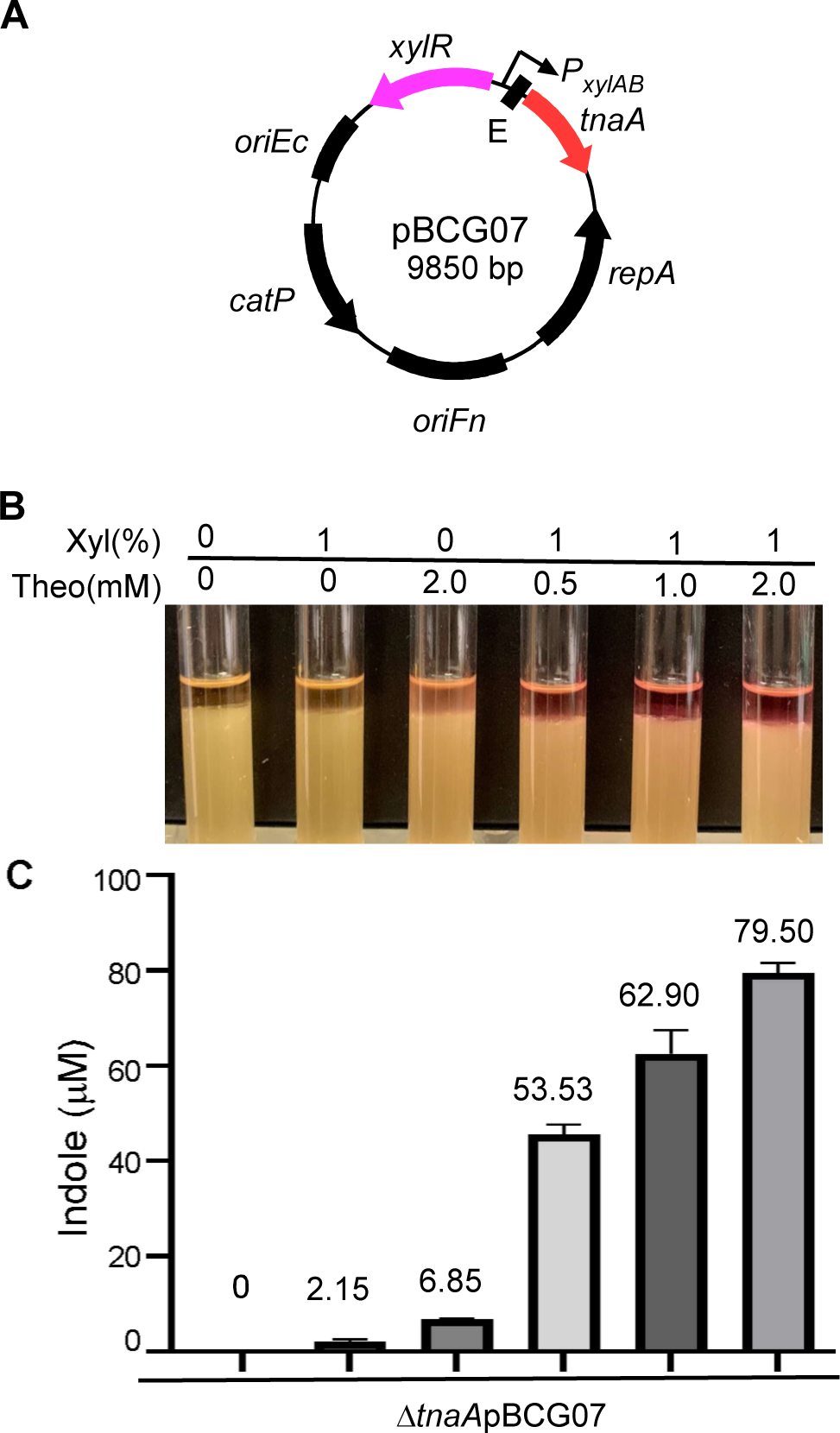
Titration of *tnaA* reporter gene expression using the dual control inducible system in *F. nucleatum*. **(A).** Genetic map of pBCG07 for gene expression studies. The expression of *tnaA*, a gene for tryptophanase that converts tryptophan to indole, is controlled by P*_xylAB_* and riboswitch at both the transcriptional and translation levels. **(B)** TnaA inducible expression level in response to the inducer(s) concentration. The TnaA activity can be monitored by indole production. The indole production was demonstrated by adding Kovac’s reagent (500 µl), which acts with the indole giving a red color in 2ml fusobacterial cultures. *F. nucleatum* Δ*tnaA* strain carrying pBCG07 was cultivated in a TSPC medium in the presence of a different concentration of inducers for 12 hours of growth. **(C)** Indole production due to TnaA presence was qualified by indole assay. 100 µl Aliquots of each culture were mixed with an equal volume of Kovac’s reagent in a 96-well plate. Tap the plate to mix briefly and thoroughly; then measure the absorbance at 565 nm. Indole levels in unknowns were calculated by comparison of absorbance values to those of a standard curve run in the same experiment. The measurement of indole production was repeated three times, and the mean values of one representative experiment performed in triplicate are reported and indicated.

### Application of this XIS-riboswitch system to study FtsX function in *F. nucleatum*

Having demonstrated that the XIS-riboswitch is a non-leaky, inducible system that allows for fine-tuning gene expression, we intend to assess further its applicability by studying the *ftsX* gene. The FtsX protein functions as a bacterial ABC transporter, controlling the activity of periplasmic peptidoglycan amidase through its interaction with the murein hydrolase activator, EnvC (48). In *F. nucleatum,* FtsX is essential for separating daughter cells following cell division. Our previous findings showed that without FtsX, cells develop into elongated filamentous structures (28).

To further investigate FtsX function in *F. nucleatum*, we created the plasmid pBCG08, which is the expression of a C-terminal Flagged *ftsX* gene under the control of the XIS-riboswitch (Figure 3A). We introduced this plasmid into an *ftsX* mutant strain, resulting in the transformant Δ*ftsX* pBCG08. Subsequently, we cultured the cells of this transformant in the presence of various combinations of the two inducers, xylose, and theophylline. In the absence of inducers, cells displayed a phenotype akin to Δ*ftsX* cells, forming elongated filaments. With 1 mM xylose alone, cells still bore a resemblance to the mutant cells; however, when 2 mM theophylline was added, cells began to shorten, implying that the xylose-inducing regulon was leaker than the riboswitch unit (Figure 3B, the top panel, the 2nd and 3rd pictures). This finding is consistent with our reporter assay data. The presence of 2 mM theophylline alone led to cell shortening, indicating the occurrence of a leak. However, the anti-FtsX antibody failed to detect the amount of leaky FtsX in the condition (Figure 3C).

**Figure 3:**
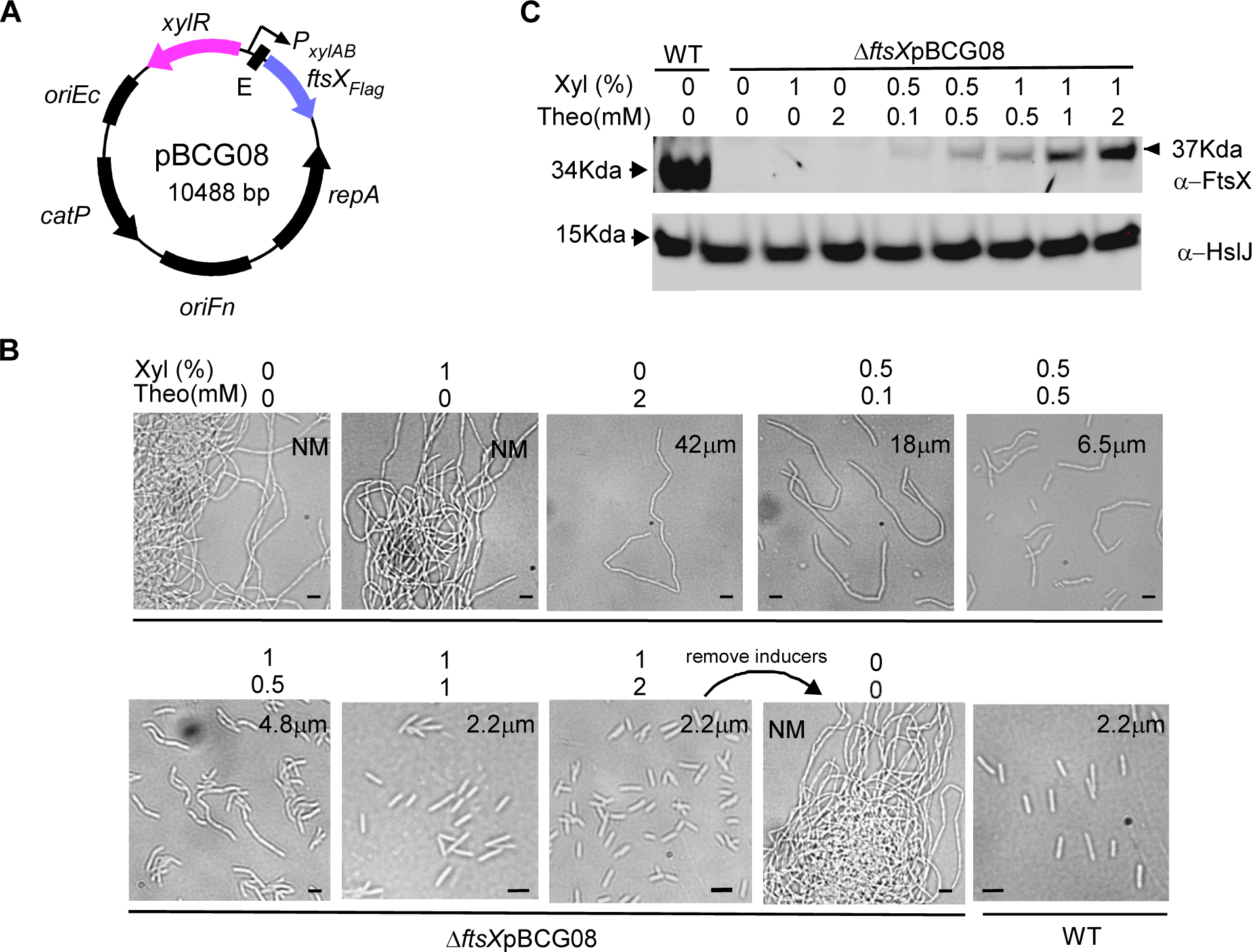
The dual inducible system controls *ftsX* expression to study its function in cell division by *F. nucleatum*. **(A)** Genetic map of pBCG08 for controlling *its* expression under the control of P*_xylAB_* and riboswitch-based regulation. **(B)** Induction of FtsX at different levels alters fusobacterial cell morphology. *F. nucleatum* Δ*ftsX* strain harboring pBCG08 was grown in a TSPC medium containing various xylose and theophylline concentrations for 10 hours of growth in the anaerobic chamber. The sample induced by 1% xylose and 1 mM theophylline after 10 hours of growth was washed twice with water to deplete the inducers and was re-inoculated into a fresh medium without an inducer for another 12 hours. A Phase-contrast microscopy recorded cell shape. NM donates instances where measurements were not obtained due to excessive cell length; Average cell lengths were computed based on 120 randomly selected observation across three distinct fields of view. Bars, 2 µm. **(C)** An equal number of cells corresponding to the panels in (B) and two additional samples from wild-type and the one induced by 1% xylose and 2 mM theophylline were subject to SDS-PAGE analysis by immunoblotting with antibodies against FtsX (α-FtsX) and HsIJ (α-HsIJ); the latter serves as a loading control. The positions of molecular mass markers (in kilodaltons) are indicated to the left of the blot. Of note, the *ftsX* gene in pBCG08 contains a 3xFlag tag.

As we raised the inducer concentrations, bacterial cell length gradually decreased. Cell length was nearly identical to wild-type cells at 1% xylose and 1 mM theophylline (Figure 3B), despite the induced FtsX level being significantly lower than in wild-type cells (Figure 3C). The western blot analysis showed increased FtsX expression levels correlated with the inducer concentrations (Figure 3C).

Lastly, we examined if gene expression could be reversed by removing the inducer. Strikingly, when inducers were removed from a culture containing 1% xylose and 2 mM theophylline, cell shape-shifted from average to elongated filamentous structures again (Figure 3B, the bottom panel, the 3rd and 4th pictures) after 8 hours, these results demonstrate that the XIS-riboswitch is an effective inducible system for precise gene expression control in *F. nucleatum*.

### Application of this XIS-riboswitch system to study *lepB*, an essential gene in *F. nucleatum*

Creating this leak-free inducible system was initially driven by our desire to investigate essential genes, as deleting them from the chromosome is only possible when a functional substitute copy exists. Maintaining rigorous control over this copy allows us to intentionally inactivate essential genes under specific conditions, potentially uncovering their functions and significance. The *lepB* gene, which encodes the type 1 signal peptidase, emerged as a potential essential gene in our experiment. LepB typically has numerous substrates, but we are particularly interested in the LepB role in the biogenesis of outer membrane protein RadD, which is crucial for dental plaque formation due to its involvement in fusobacterial physical interactions with other bacteria. The RadD protein precursor contains an N-terminal signal peptide, which is presumably cleaved by the type I signal peptidase upon transport to the inner membrane. This allows for proper RadD protein folding and secretion, ultimately displaying on the outer membrane. We hypothesized that without LepB, RadD levels would decline or vanish entirely.

To examine this hypothesis, we endeavored to construct a *lepB* in-frame deletion strain of *F. nucleatum* using a two-step recombination method we had previously developed (28). We cloned approximately 1.7-kilobase DNA fragments flanking *lepB* into the non-replicating vector pCM-*galK*, which produces expressing galactokinase (GalK) as a counter-selectable marker. After transforming the resultant plasmid for allelic exchange into a Δ*galK* strain, we produced single-crossover strains via homologous recombination. A double crossover event in this strain could replace the wild-type gene with the deletion copy or a reversion to the wild-type. After isolating the double-crossover strains through *galK* counterselection, we tested 69 2-deoxy-D-galactose (2-DG)-resistant colonies and found none were mutants, and all isolates carried the wild-type allele. This outcome suggested that the *lepB* is essential for growth in *F. nucleatum*.

To confirm this, we sought to isolate a *lepB* mutant strain in a *lepB*/Δ*lepB* merodiploid background with a functional *lepB* copy in a shuttle plasmid pCWU6, controlled under by the XIS-riboswitch inducible system. The pCWU6 plasmid contains a chloramphenicol resistance marker (*catP*), the same as the maker in the in-frame deletion plasmid pCM-*galK*. To ensure compatibility of antibiotic selection between the in-frame deletion plasmid and pCWU6-based expression vector, we first created a suicide plasmid, pBCG10, where the erythromycin resistance cassette *ermF-ermAM* replaced *catP* in pCM-*galK* and made a new *lepB* in-frame deletion plasmid pBCG10-Δ*lepB* (Figure 4A). Utilizing pBCG10-Δ*lepB*, we re-obtained the *lepB*/Δ*lepB* merodiploid strain via single crossover (refer to steps 1 and 2 in Figure 4B). We subsequently created pAN01, in which the XIS-riboswitch system regulates *lepB* expression. The pAN01 was transformed into the *lepB*/Δ*lepB* merodiploid strain (refer to step 3 in Figure 4B), and the resulting transformants were cultured and plated on plates containing 2-deoxy-D-galactose. This metabolite becomes toxic as 2-deoxygalactose-1-phosphate in cells that retain pBCG10 due to *galK*. The selected plates also included the inducers (1% xylose and 2 mM theophylline) to induce *lepB* expression within the cells. In this genetic background, 6 of 11 double-crossover strains carried the deletion allele (see step 4 in Figure 4B and Figure 4C). In contrast, when the plates lacked inducers, all isolates were the wild-type allele. These results demonstrated that *lepB* is essential under the condition tested.

**Figure 4:**
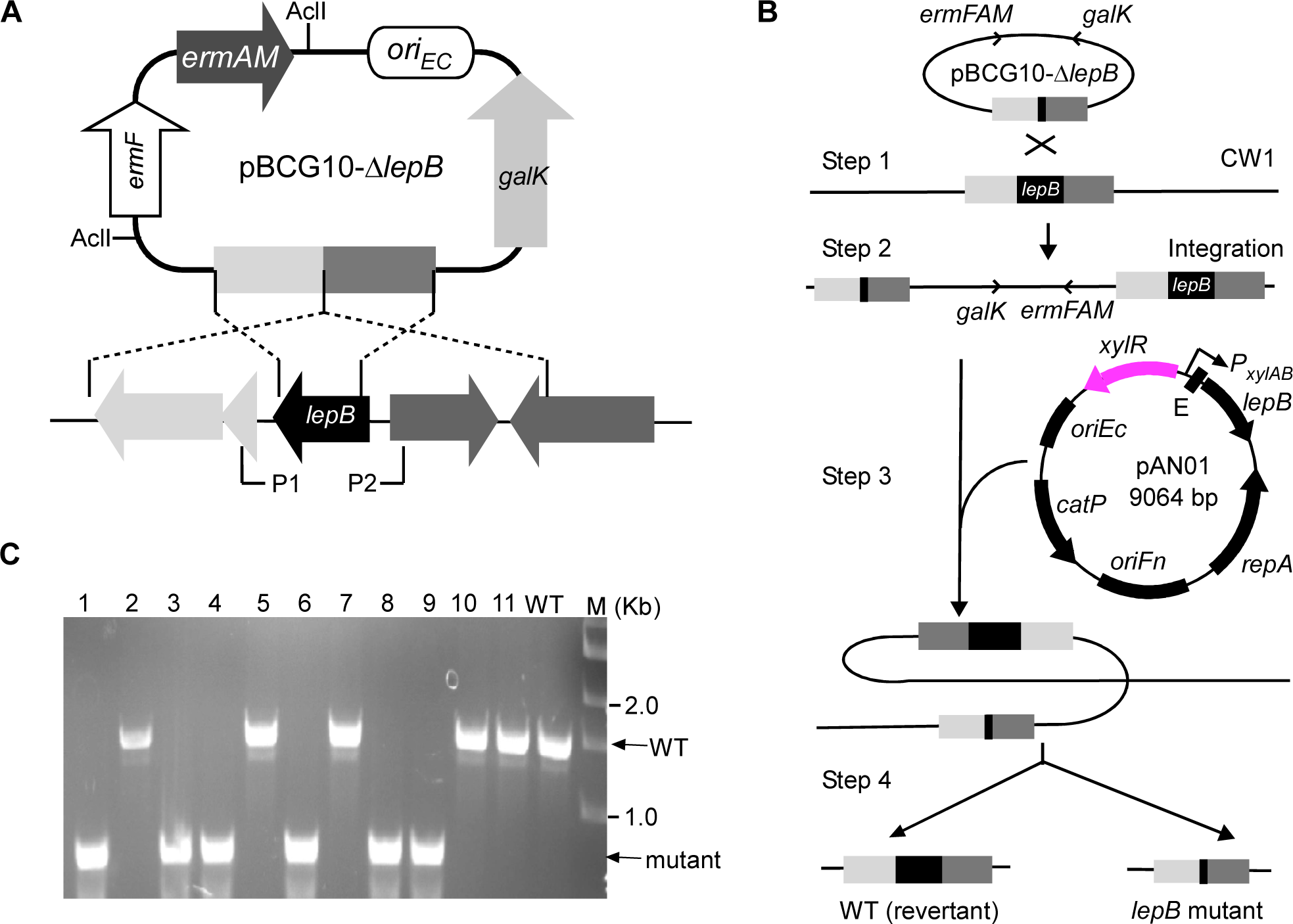
The use of the dual inducible system to make a deletion of the type I signal peptidase *lepB* in *F. nucleatum*. **(A)** Presented here is the non-replicative vector pBCG10 used to produce unmarked, in-frame deletions in *F. nucleatum*. The vector carries a 1.7 kb flanking region of *lepB*, the erythromycin resistance cassette (*ermF-AM*), and *galK*. Brackets indicate regions amplified by a primer pair P1 and P2. **(B)** The schematic diagram is shown *lepB* allelic exchange with pBCG10-Δ*lepB* via a two-step strategy. Integration of this vector into the *F. nucleatum* Δ*galK* mutant CW1 generates a *lepB*/Δ*lepB* merodiploid strain (single-crossover strain). Excision of *lepB* from integrated strain chromosome is permitted in the presence of pAN01, a plasmid expressing a functional copy of the *lepB* gene under the control of a xylose-inducible promoter in combination with a theophylline-responsive riboswitch E element, resulting in *lepB* in-frame deletion mutant and wild-type alleles (double-crossover strains). **(C)** The excision of l*epB* from the chromosome of mutant strains was confirmed by PCR amplification using the primer pair P1 and P2.

To demonstrate that *lepB* is required for cell viability, we compared the growth rates of the *lepB* conditional mutant in broth media supplemented with various inducers of xylose and theophylline concentrations. We used the wild-type strain containing an empty vector as a control. The growth of the *lepB* conditional mutant strain was comparable to the wild-type strain when 0.5% xylose and 1 mM theophylline were present. However, when inducer concentrations were lowered, growth was notably inhibited. The *lepB* mutant did not grow without inducers (Figure 5A). Simultaneously, we examined serially diluted aliquots from the same set of cultures on agar plates, with or without xylose and theophylline, alone or in combination. We observed a growth pattern similar to the *lepB* broth’s growth. Furthermore, we noted that cell growth with only 2 mM theophylline was weaker than when cultured with 0.1% xylose and 0.2 mM theophylline but better than with just 1% xylose added (Figure 5B). These experimental results demonstrated that *lepB* expression is tightly regulated by the XIS-riboswitch system, and the growth of the *lepB* conditional mutant directly depends on the presence of inducers. LepB mutant cells can grow when inducers are present, but not in their absence underlines the essential role of *lepB* in *F. nucleatum*’s viability.

**Figure 5.**
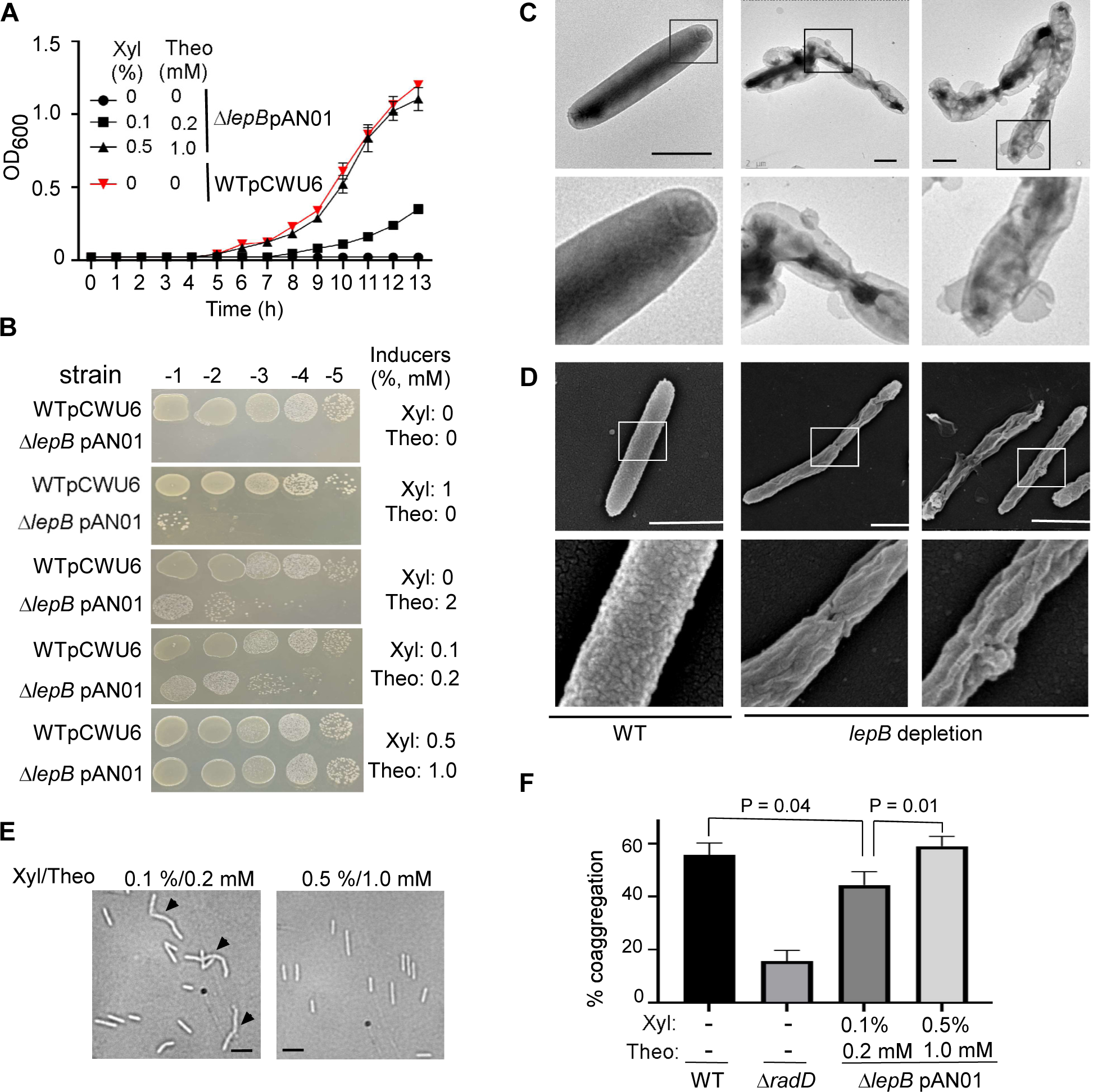
The type I signal peptidase *lepB* in *F. nucleatum* is an essential gene. **(A)** The effect of inducer(s) on the growth of conditional *lepB* mutants in the liquid media. *F. nucleatum* strains (wild-type: strain ATCC 23726 harboring an empty vector pCWU6 (red triangle); *lepB* conditional mutant: Δ*lepB* pAN01) were grown statically under anaerobic conditions on TSPC media with various doses of inducers (xylose and theophylline). Turbidity was monitored at an optical density of 600 nm. Data represented the means of standard deviation of the results of triplicate biological experiments. **(B)** The effects of inducer(s) on the growth of conditional *lepB* mutants in solid media. Ten-fold serial dilutions of overnight cultures of the wild-type 23726 and the conditional Δ*lepB* strains were spotted on agar plates with and without inducer(s) xylose and theophylline and cultivated in an anaerobic chamber at 37°C. Cell growth was recorded after three days of incubation. This experiment was repeated three times, and one representative experiment was shown. **(C)** LepB depletion creates short filaments and alters outer membrane structures. The *lepB-depleted* and lepB-inducing cells were immobilized on carbon-coated nickel grids and stained with 0.1% uranyl acetate before viewing with a transmission electron microscope. Enlarged areas of the top panels are shown in the bottom panels. Bars, 1 µm. **(D)** The altered cell surface of conditional *lepB* mutant was revealed by scanning electron microscopy. The cells from *lepB* depletion or *lepB* induction were immobilized on a silicon wafer and fixed with 2.5% glutaraldehyde before viewing by a scanning electron microscope. Enlarge areas of the top panel are shown in the below panels. Bars, 1 µm. **(E)** Comparison of lepB conditional mutant strain umorphology. Cells were grown under two distinct conditions with inducers (0.1% xylose/0.2 mM theophylline and 0.5% xylose/1.0 mM theophylline). Microscopic examination was conducted after cells reached the stationary phase to assess morphological differences. **(F)** The quantitative coaggregation assay was performed by mixing wild-type *F. nucleatum*, *radD*, and *lepB* mutants with an equal number of *A. oris* cells in the coaggregation buffer (CAB). The mixture was allowed to aggregate for 20 mins before measuring the OD_600_ absorbance. Measurements were taken both before and after the aggregation process. The presented data represent the average of three independent experiments. Statistical analysis was performed using Student’s *t*-test with GraphPad Prism software.

To understand why the lack of *lepB* expression results in the lethality of *F. nucleatum*, we first examined the surface structure of the conditional *lepB* mutant cells under conditions without inducers. The *lepB* mutant cells were initially cultured in a medium containing 0.5% xylose and 1 mM theophylline. During the stationary phase, cells were collected and washed three times with a pre-reduced medium to eliminate residual inducers. Then, cells were plated on plates without inducers and incubated for 16 hours. Upon harvesting, the cells were stained with uranyl acetate for negative staining and observed under an electron microscope. Interestingly, the mutant cells displayed significant morphological abnormalities. They had an unusual outer membrane structure with vesicle-like formation on the cell surface and formed short chains of 2-3 cells. These cells were further analyzed using a scanning electron microscope after fixation with glutaraldehyde and alcohol dehydration. Compared to the wild-type cell surfaces, which have a uniformly distributed protruding structure similar to a knot in a carpet, the *lepB* cells without inducers exhibited uneven surfaces with hemp rope-like structures. The evidence showed that *lepB* gene expression is crucial for outer membrane integrity and cell division.

We further investigated if LepB plays a role in presenting fusobacterial RadD on the cell surface. However, when grown without inducers, Δ*lepB* cells become dying and prone to autoaggregation, making them unsuitable for coaggregation assays with its partners. To overcome this, we employed Δl*epB* cells grown in two distinct inducer concentrations (0.1% xylose/0.2 mM theophylline vs. 0.5% xylose/1.0 mM theophylline). Cells from both conditions did not aggregate, although some formed short chains in the lower inducer medium (Figure 5E). As inducer concentrations influence *lepB* expression levels, we expected cells grown in a high-inducer medium to produce more LepB. This led to increased RadD delivery to the surface and improved coaggregation with *Actinomyces oris* . This hypothesis was confirmed when cells from the high-inducer medium exhibited a higher coaggregation rate than those from the lower inducers. This evidence indicates that in *F. nucleatum*, the RadD precursor is a substrate for LepB, emphasizing the importance of LepB in RadD presentation on the cell surface.

## DISCUSSION

A tightly regulated inducible system is a powerful tool for studying gene function, especially for essential genes. In this study, we combined the xylose-inducible system with theophylline-responsible riboswitch. We created a hybrid inducible system that controls the expression of our interested genes at both transcriptional initiation and translation level. Using two reporters, the luciferase and indole-producing tryptophanase, we showed that this combination inducible system is leak-free. In the absence of inducers, no or negligible signal was detected. We used this system to fine-tune the expression of FtsX and elegantly showed a negative correlation between FtsX level and cell length (Figure 3). Most importantly, this leak-free system can study *lepB*, an essential gene in *F. nucleatum*.

An essential gene is a gene that is critical for the survival and reproduction of an organism under normal growth conditions. The loss or inactivation of an essential gene typically results in lethality or severely compromises the growth and viability of the bacteria. Essential genes are of particular interest to researchers, as they often encode proteins involved in fundamental cellular processes and may represent targets for developing new antimicrobial drugs (49). Bacterial type I signal peptidase (SPase I) is essential to the Tat and Sec secretory systems. SPase I is responsible for the hydrolysis of the N-terminal signal peptides from proteins secreted across the cytoplasmic membrane and plays a key role in bacterial viability and virulence (24). Thus, type I SPase is considered to be an attractive target (50, 51). *F. nucleatum* possesses a LepB homolog, and a recent study revealed by RNA-seq that *lepB* is controlled by σ ^E^, a mediator of an oxygen-induced response (27). However, no genetic study of *lepB* in *F. nucleatum* has been conducted. In our experiments, we pursued an answer to whether LepB plays a role in the cell’s presentation of fusobacterial adhesin RadD. We attempted to construct Δ*lepB* mutants using a *galk*-based two-step strategy but were unsuccessful. However, we could delete the chromosomal copy only when a second functional copy was provided elsewhere (Figure 4B). By placing expression under the control of the leak-free XIS-riboswitch regulon (Figure 4A and 4B), we confirmed that depletion of *lepB* expression was detrimental to growth in the absence of inducers (Figure 5A and 5B). Further, without *lepB*, we revealed that the bacterial outer membrane structure underwent significant changes, and cell division could not proceed well (Figures 5C and 5D). By controlling *lepB* expression by regulating the addition of two inducers, we discovered that the amount of *lepB* expression directly influences fusobacterial coaggregation with its partner *A. oris*, possibly due to reduced RadD surface display when LepB was limited. RadD mediates *F. nucleatum* aggregation with *A. oris*.

Studying the *lepB* essential gene benefited from constructing this leak-free inducible system. Currently, there are two gene expression inducible systems used in *F. nucleatum*. One is the tetracycline-inducible (Tet) system (27), and the other is riboswitch-based (32). The two systems and other most inducible systems used in the bacterial genetic study have some degree of leakage (34, 52). To address this issue, researchers have employed various strategies. Some common approaches involve increasing the expression of repressor proteins through stronger constitutive promoters or adding more operator sequences to minimize leak levels. Another method involves combining two different inducible systems. The inducible systems are mainly divided into two categories: those regulated at the transcriptional level, like the Tet system and XIS, and those that control translation initiation, like the riboswitches. By integrating these two types of systems, researchers have been able to construct a dual transcriptional-translation control offers exceptionally low leakage expression. Our laboratory has pioneered this approach by combining the Tet system with a riboswitch unit. We had successfully applied this novel system to study essential genes in the oral bacterium *A. oris* (29). Following our success, other researchers have developed similar combined systems (52, 53). Combining the riboswitch with the xylose-inducible system has yet to be previously attempted; we are the first to demonstrate that this combination system can suppress entirely leakage transcription in *F. nucleatum*. This system is not only applicable for studying gene expression in *F. nucleatum* subsp. *nucleatum* but also in other subspecies strains (Figure S2). In comparison to its expression in subsp. *nucleatum*, luciferase activity in subsp. *animalis* is lower under the same induction conditions. This discrepancy could be attributed to differences in inducer permeability between the two bacterial types or an additional repression regulatory element in subsp. *animalis* that requires further investigation. However, in the absence of inducers, this system exhibits no leakage, suggesting that it may be suitable for use in other subspecies of *F. nucleatum*.

The luciferase gene, widely used as a gene reporter in various bacterial genetic studies, requires oxygen to detect its activity (17, 54, 55). *F. nucleatum* is an anaerobic bacterium; therefore, we must remove fusobacterial cultures from an anaerobic environment to measure luciferase activity and expose them to oxygen. When vigorously mixed with a pipettor multiple times, we found that a small volume of culture, such as 100 µl, can obtain sufficient oxygen to support the luciferase reaction. Recently, GFP and mCherry fluorescent proteins have been used as reporters in *F. nucleatum* studies (27). Nonetheless, the maturation of GFP and mCherry necessitates a complex chemical fixation process and a lengthy incubation in a cold environment (27). In comparison, luciferase is a more convenient reporter if used solely to monitor changes in gene expression. However, GFP and mCherry possess a unique advantage, as they can be used for studying protein localization when translationally fused to the target gene. Both luciferase and fluorescent proteins require complex and expensive instruments for detection. An ideal promoter reporter that could be easily observed with the naked eye would involve simple experiments, which led us to consider indole and the indole-producing enzyme tryptophanase gene *tnaA*. Indole is easily detected, and its color intensity allows for a rough estimation of gene promoter strength. By placing this gene under the control of the XIS-riboswitch, we observed no red color without inducers; as inducers were gradually added, the red color intensified. Furthermore, the produced indole can be quantitatively detected. Given the ease of indole detection and the ability to estimate its production based on color intensity, the *tnaA* gene as a reporter could potentially be employed for large-scale screening in bacterial genetic studies. By transcriptionally fusing to the target gene and incorporating Tn5 transposition, it can aid in the screening of factors influencing the expression of the target gene. As *F. nucleatum* naturally possesses a *tnaA* gene, the downside is that these experiments must be performed in a *tnaA* mutant background.

In summary, we have successfully developed a leak-free inducible gene expression system in *F. nucleatum*, enabling us to investigate *ftsX* and *lepB* (an essential gene) functions. By devising this hybrid system, we have broadened the range of genetic tools available for *F. nucleatum* research. This advancement will assist in the dissection and comprehension of the roles of other vital and essential genes within this microorganism. As a result, our work could enhance the understanding of *F. nucleatum* biology. It may pave the way for the development of innovative strategies to prevent and treat diseases associated with *F. nucleatum*. In addition to our work with *F. nucleatum*, the leak-free inducible system could be adapted and applied to other bacterial species, further expanding its utility in microbial genetics.

## MATERIALS AND METHODS

### Bacterial strains, media, and growth conditions

Bacterial strains are listed in Table 1. *F. nucleatum* strains were grown in a TSPC medium consisting of 3% tryptic soy broth (TSB, BD), 1% Bacto^TM^ peptone, and 0.05% cysteine added prior to inoculation or on TSPC agar plates in an anaerobic chamber filled with 80% N_2_, 10% H_2,_ and 10% CO_2_. *Escherichia coli* strains were grown in Luria-Bertani (LB) broth. *F. nucleatum* strains harboring pBCG06, pBCG06a, pBCG07, pBCG08, or pAN01 were grown with 5 µg/ml thiamphenicol in the presence of various dose combinations of two inducers (xylose and theophylline). The 40 mM theophylline and 20% xylose stocks independently were prepared in a TSP medium, kept at -4°C for long-time storage, and warmed to 37°C before use. When required, Antibiotics used as needed were: kanamycin (50 µg ml^-1^), chloramphenicol (15 µg ml^-1^), thiamphenicol (5 µg ml^-1^), erythromycin (100 µg ml^-1^) and clindamycin (1 µg ml^-1^). Reagents were purchased from Sigma unless indicated otherwise.

**Table 1:**
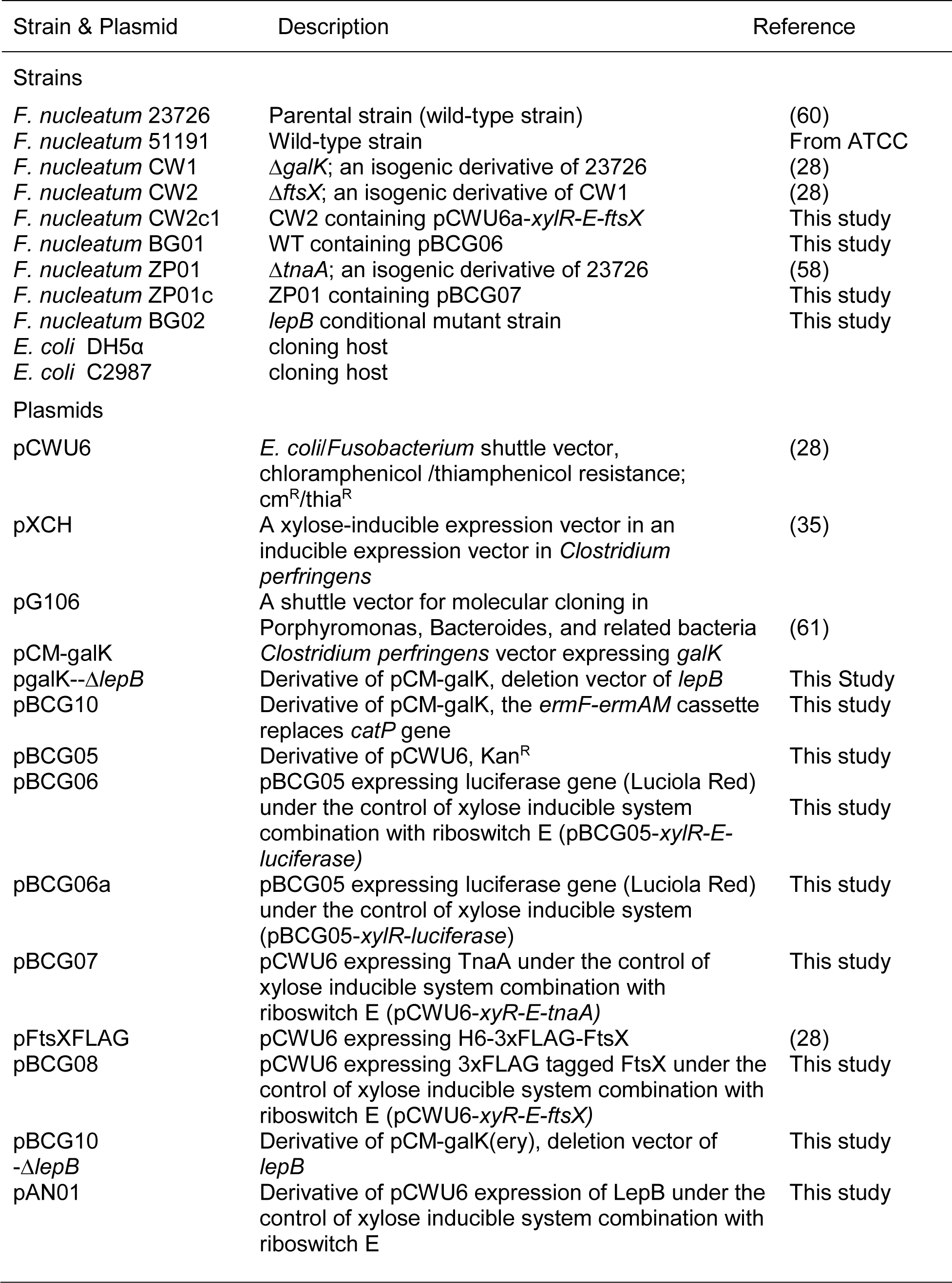
Bacterial strains and plasmids used

### Plasmid and strain construction

All plasmids are listed in Table 1, and most of them were constructed by Gibson assembly using reagents from New England Biolabs, following the manufacturers’ instructions. In brief, a 50-100 ng linearized cloning vector obtained from a restriction enzyme digestion or an inverse PCR was mixed with 150-400 ng inserted PCR fragments with 10 µl 2x Gibson Assembly Master Mix in a final volume of 20 µl at 50°C for 20-60 minutes. The *E. coli* competent cells were transformed with 2-5 µl of the master mix/vector/inserts. Regions of plasmids constructed using PCR were verified by DNA sequencing. The oligonucleotide primers used in this work (Table 2) were synthesized by Sigma Aldric. All plasmids were propagated using *E. coli* DH5a or C2987 as the cloning hosts and introduced into fusobacterial strains by electroporation. All inducible plasmids were built on the backbone of pCWU6 (28), an *E. Coli*/*Fnucleatum* shuttle vector with a chloramphenicol resistance marker.

**Table 2:**
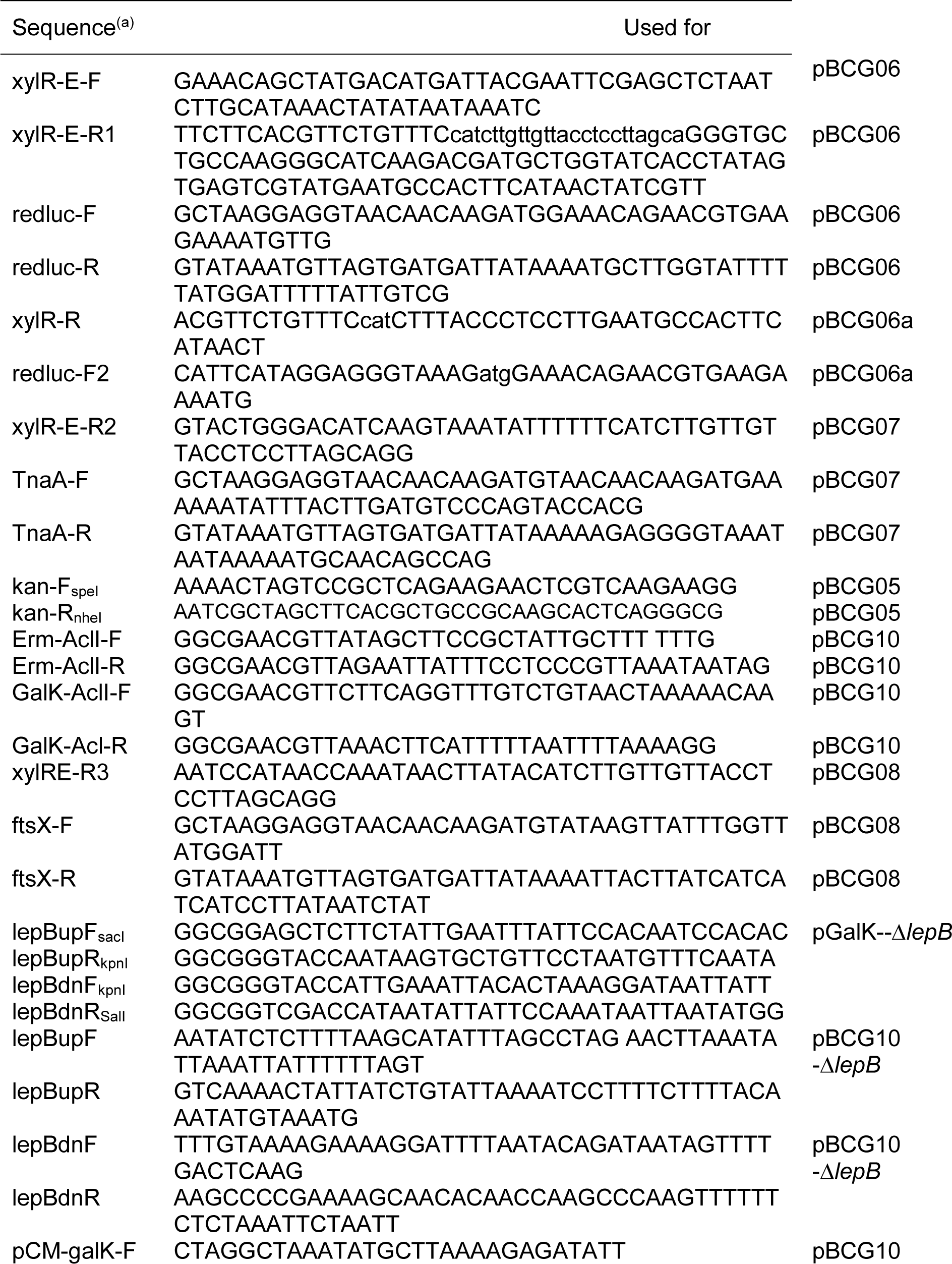
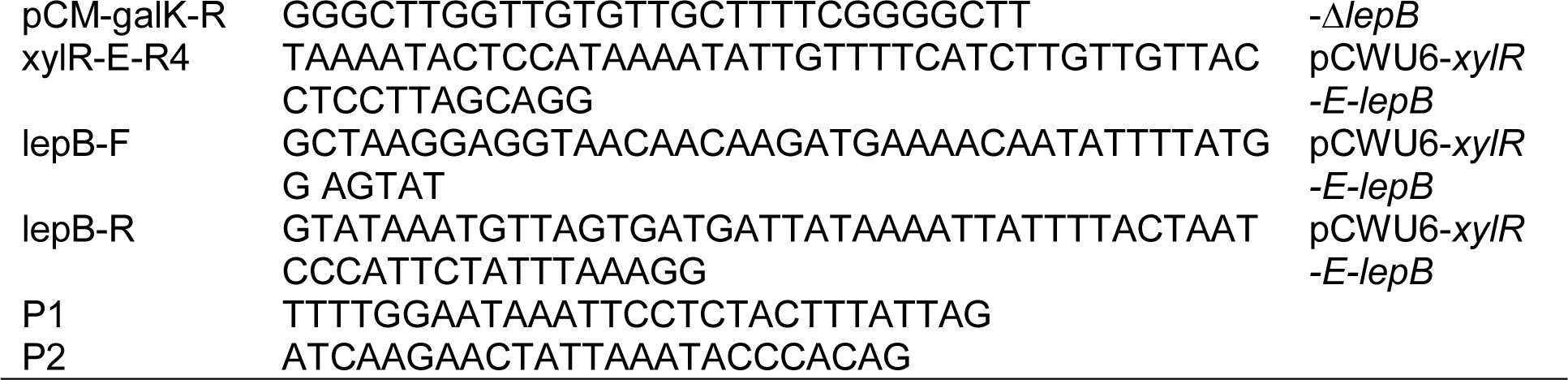
Primers used in this study

pBCG06a-In order to assess the leak of the xylose-inducible system, we constructed this plasmid in which the reporter luciferase gene’s expression is controlled by a xylose-inducible promoter alone. PCR obtained the *xylR*::P*_xylAB_* fragment with the plasmid pXCH (35) as a DNA template. The primer pair redluc-F2/redluc-R was used to PCR amplify the luciola red luciferase gene (*luc*) from *S. mutans* strain JK-G-3 (55). The two amplicons of *xylR*::P*_xylAB_* and *luc* gene were fused by an overlapping PCR. The resultant product was mixed in the Gibson assembly Master mix solution with SacI/HindIII-cut pBCG05 to generate pBCG06a. pBCG05 is a derivative vector of pCWU6, harbors a kanamycin (*kan*) resistant gene cassette from pJRD215 (56) inserted at sites SpeI and NheI of pCWU6. pBCG05 allowed *E. coli* to grow faster on LB plates with 50 µg/ml kanamycin than with 15 µg/ml chloramphenicol, shortening the screening time for positive clones.

pBCG06-To prevent the leak of the xylose-inducible system, a theophylline-responsible riboswitch E unit (34) was incorporated into the 5’ untranslated region (5’ UTR) of the reporter *luc* gene to replace the original ribosome binding site (RBS) and the short spacer sequence in between RBS and the start codon in pBCG06a (see Figure 1B and 1D). The riboswitch E sequence was added at the 5’ end of the primer xylR-E-R1 to do it. This primer works with xylR-E-F, obtaining *xylR*::P*_xylAB_*::E by PCR amplification from pXCH. The pBCG06a was used as a DNA template to amplify the *luc* gene with primers redluc-F/R. An overlapping PCR reaction was used to fuse the two amplicons *xylR*::P*_xylAB_*::E and *luc* gene. The fusion product was cloned into SacI and HindIII-digested pBCG05 by Gibson assembly and transformed into *E—coli* C2987 cells, as described previously (57).

pBCG07-To test if the indole produced by the tryptophanase (TnaA) works as a reporter in *F. nucleatum* for gene expression, we PCR-amplified *xylR*::P*_xylAB_*::E DNA fragment from pBCG06 with primer pair xylR-E-F/R2.The primer pair TnaA-F/R were used to amplify the *tnaA* coding region from the genomic DNA of *F. nucleatum* ATCC 23726. The two PCR products were mixed and cloned into SacI and HindIII-digested pCWU6 by Gibson assembly. DNA sequencing further confirmed the resultant plasmid and subsequently electroporated into the Δ*tnaA* mutant (58).

pBCG08-To validate the utility of the xylose and theophylline duel control system for deciphering gene function in *F. nucleatum*, we induced the expression of *ftsX*, a gene involved in cell division, in the presence of both inducers at different doses. To construct this vector, primer sets xylR-E-F/R3 and ftsX-F/R were used to amplify the *xylR*::P*_xylAB_*::E from pBCG06 and the *ftsX* coding region, including a 3x Flag tag from the plasmid pFtsX_Flag_ (28). The two PCR products were ligated into SacI and HindIII-digested pCWU6 by Gibson assembly.

pAN01-To further test the dual inducible system’s use in studying the essential genes in *F. nucleatum*, we PCR-amplified *xylR*::P*_xylAB_*::E DNA fragment from pBCG06 with primer pair xylR-E-F/R3. The *lepB* gene was amplified from the genomic DNA of *F. nucleatum* ATCC 23726 with primers lepB-F/R. The *xylR*::P*_xylAB_*::E was cloned upstream of *lepB* via Gibson assembly to create *xylR*::P*_xylAB_*::E:*lepB* inducible plasmid named pAN01.

pGalK-Δ*lepB-* To create deletion constructs of *lepB*, appropriately 1.7-kb fragments upstream and downstream of *lepB* were amplified by PCR using lepBupF_sacI_/lepBupF_KpnI_, lepBdnF_KpnI_/lpeBdnR_salI_. The upstream fragments were digested with SacI and KpnI, while the downstream fragments were digested with KpnI and SalI. These fragments were ligated into pCM-*galK* precut with SacI and SalI. The generated plasmid was confirmed by restriction enzyme digestion.

pBCG10-Δ*lepB*-pCM-galK containing a chloramphenicol resistance marker (*catP*) is the suicide plasmid used for gene deletion in *F. nucleatum*. To work with pCWU6-based plasmid pAN01, the *catP* in pCM-galK was replaced with the erythromycin resistance cassette *ermF-ermAM* to create pBCG10. Primers erm-Acll-F and erm-Acll-R, each containing an Acll site for cloning, were used along with pG106 as a template to PCR-amplify a DNA segment encompassing the complete erythromycin resistance cassette *ermF-ermAM*. PCR linearized pCM-galK with primer set GalK-Acll-F/R to remove the *catP* gene. The linearized pCM-galK backbone and *ermF-ermAM* were

digested by Acll and ligated together to generate pBCG10. To create an in-frame deletion construct of *lepB*, 1.0 kb fragments upstream and downstream of *lepB* were amplified by PCR using lepBupF/R and lepBdnF/R primers. An overlapping PCR reaction fused the upstream and downstream fragments. The primer pair pCM-galK-F/R was used to amplify the backbone of pBCG10. The pBCG10 vector backbone ligated with the fused upstream and downstream of *lepB* via Gibson assembly to create pBCG10--Δ*lepB*. pBCG10-Δ*lepB* were further confirmed by DNA sequencing and transformed into *F. nucleatum* CW1 strain by electroporation.

All produced plasmids were confirmed by sequencing and then transformed into *F. nucleatum* ATCC 23726 or *F*. *nucleatum* ATCC 51191 with the electroporation procedure we previously published (59). In brief, cells of either ATCC 23726 or ATCC 51191 were harvested by centrifugation from a 100 ml stationary phase culture. The cells were washed twice with sterile water and once with 10% glycerol, then resuspended in 3 ml of 10% glycerol and divided into 0.2 ml aliquots. Next, 1.0 µg purified plasmid DNA was added to 0.2 ml aliquot of the above-prepared fusobacterial cells in a pre-cooled corvette (0.1 cm electrode gap, Bio-Rad) and left on ice for 10 minutes. The reactions were electroporated using a Bio-Rad Gene Pulser II set at 25 kV, 25 µF, and 200 Ω. Immediately following electroporation, the cells were diluted in 1 ml of pre-reduced and pre-warmed TSPC. The culture was plated on TSPC agar plates with 5 g/ml thiamphenicol after being incubated anaerobically for 5 hours without agitation.

### Initial attempts to make an in-frame deletion of *lepB*

A published procedure based on GalK as a counterselection marker (28) was followed to generate non-polar, in-frame deletion mutants in *F. nucleatum*. Briefly, the *lepB* deletion plasmid pGalK-Δ*lepB* or pBCG10-Δ*lepB* was electroporated into CW1, an isogenic mutant of *F. nucleatum* 23726 that lacks *galK* (28). Integrating the vector into the bacterial chromosome via homologous recombination was selected on TSPC agar plates containing 5 µg ml^-1^ thiamphenicol or 1 µg ml^-1^ clindamycin. Excision of the vector via the second recombination event – which results in possible gene deletion or reconstitution of the wild-type genotype – was selected by 0.25% 2-deoxy-D-galactose (2-DG). 2-DG-resistant and thiamphenicol-sensitive colonies were screened for the expected deletion mutation by PCR amplification. We PCR-screened about 100 colonies (69 colonies for pGalK-Δ*lepB* and 31 for pBCG10-Δ*lepB*). Of 100 colonies tested, all possessed a WT genotype, and no in-frame deletion strains were isolated. This suggests that *lepB* is an essential gene in the condition we tested.

### Creation of a conditional *lepB* deletion mutant

To delete the *lepB* gene from the chromosome, the strain with pBCG10-Δ*lepB* (Integrant cells) was then transformed with pAN01 expression, a functional copy of the *lepB* gene under control by the xylose-inducible promoter and riboswitch-based unit. The resultant transformants were selected on TSPC agar plates supplemented with 5 µg ml^-1^ thiamphenicol and 1 µg ml^-1^ clindamycin. A colony of transformants was used to inoculate an overnight culture in TSPC (with thiamphenicol alone) containing two inducers (1% xylose and 2 mM theophylline). 100-µl aliquots of 100-fold diluted overnight cultures in fresh TSPC were spread out on TSPC agar plates containing 0.25% 2-deoxy-D-galactose (2-DG), thiamphenicol, and both inducers. After three days of incubation at 37°C in an anaerobic chamber, the grown colonies were screened for their clindamycin sensitivity. Clindamycin -sensitive colonies were then screened for the loss of the *lepB* chromosomal copy by PCR amplification. The generated conditional deletion mutant of *lepB* was analyzed for cell growth in broth or solid plates.

### Luciferase assays

An overnight culture of wild-type strain harboring the plasmid pBCG06 was washed in TSPC and diluted 1:20 in a fresh TSPC medium. The cultures were grown with 5 µg ml^-1^ thiamphenicol at 37°C in an anaerobic chamber. When the culture turbidity reached OD600 ∼0.65, the inducers were added with various dose combinations, as indicated (Figure 1F). After 2 hours of growth, three 100 μl aliquots of each culture were taken out in a 96-well microplate (#3917, Corning®). Each was mixed with 25 μl of 1 mM D-luciferin (Molecular Probe) suspended in 100 mM citrate buffer (pH 6.0), and the mixture was vigorously pipetted 15 times aerobically. Luciferase activity was measured by using a GloMax® Navigator Microplate Luminometer. Culture growth was measured spectrophotometrically by determining optical density at 600 nm. The wild-type strain with empty plasmid pCWU6 was a negative control showing a very low background noise. The cells expressing pBCG06a were used to demonstrate the leakage issue inherited in the xylose-inducible system in the absence of inducers. To test if the glucose affects the co-inducible system in *F. nucleatum*, 20 mM glucose was added to a sample with 2% xylose and 2 mM theophylline. Each experiment was repeated three times.

### Indole assay

Fusobacterial **Δ***tnaA* deletion strain containing pBCG07 was grown overnight with 5 µg ml-1 thiamphenicol and no inducers. The overnight culture was diluted 1:20 in 2ml fresh TSPC medium with the two inducers (xylose and theophylline) at different combinations. The cultures were grown statically at 37°C in anaerobic conditions. After incubation at 37°C for 15 hours, 500 µl of Kovacs’s reagent was added to each tube without shaking. A positive indole test is indicated by forming a pink-to red-color in the reagent layer on the top of the medium within 2 minutes of adding the reagent. A negative result appears yellow.

The indole assay kit (Sigma, #MAK326) was used to qualify the induction level of indole in the different inducer conditions. For the indole assays, 1 ml of each culture was centrifuged at 15,000 rpm for 2 min, and 100 µl of the supernatant was tested in triplicate. 100 µl of 4 indole standards (0, 25, 50, 100 µM) or samples of unknown indole concentrations were incubated with 100 µl Kovács reagent for 20 min at room temperature. The reaction produced a soluble product measured spectrophotometrically at 565 nm with a CYTATION 5 image reader (BioTeK). The known indole concentrations from 0 to 100 µM were tested in triplicate, and the mean results were used to construct a standard curve. Indole levels in unknown were calculated by comparison of absorbance values to those of a standard curve in the same experiment.

### Microscopic analysis on cells of Δ*ftsX* expressing various levels of FtsX

An overnight culture of **Δ***ftsX*pBCG08 (grown in TSPC medium with 5 µg ml^-1^ thiamphenicol) was sub-cultured by 1: 20 dilution in fresh TSPC medium containing various concentrations of xylose and/or theophylline as indicated (Figure 3B). The cultures were grown at 37°C for 10 hours. The cells without or with very low inducers were sedimented at the bottom of the culture glass due to forming long filaments. Thus, a strong vortex was required to obtain homogenous cell suspensions. A 2 µl of cell suspension for each culture was smeared on a glass slide, and cells were viewed using phase-contrast microscopy at 1,000x magnification.

### Western blotting analysis

If xylose and theophylline did not have an impact on bacterial growth in our experiment, then an equal volume of cultures treated with different inducers would theoretically contain an equal number of cells. 1 ml of the above-mentioned cell suspension for each culture **Δ***ftsX*pBCG08 treated with varying doses of the two inducers was used and centrifuged to collect cells. The cell pellets were washed twice with water and suspended in a sodium dodecyl sulfate (SDS) sample buffer. Samples were boiled for 10 min, and proteins were resolved on home-made 12 % SDS-polyacrylamide gel electrophoresis gel, transferred to a PVDF membrane, and probed with rabbit anti-FtsX with a 1:1000 dilution and anti-HslJ with a 1:5000 dilution as we used before (28). HslJ, a predicted inner membrane protein (HMPREF0397_1592) used as a control. Polyclonal goat anti-rabbit IgG coupled to IRDye 680LT (#92668021; LI-COR Biosciences, USA) was used to detect primary rabbit antibodies at the lowest suggested dilution of 1:5,000.

### Cell growth assay

Cells of the conditional *lepB* deletion mutant and parental strain ATCC23726 with the empty plasmid pCWU6 were grown overnight in TSPC medium containing xylose (1%), theophylline (1 mM), and thiamphenicol (5 µg ml^-1^) at 37°C in an anaerobic chamber. The overnight culture of the conditional mutant was washed three times with water to remove the two inducers and used to inoculate three cultures in fresh TSPC (1:100 dilution), each with three triplicates containing thiamphenicol and various concertation of xylose (0. 0.1, 0.5%) combination with theophylline (0, 0.2, 1.0 mM). Cell growth at 37°C was monitored by measuring optical density at 600 nm (OD_600_) every hour for 13 hours with a portable Implen^TM^ OD600 DiluPhotometer in an anaerobic chamber. The experiment was repeated twice.

For cell growth on plates, washed cells from overnight cultures were normalized to the same OD600, making 10-fold serial dilutions and spotting 8 µl on TSPC agar plates containing various concentrations of xylose (0, 0.1,0.5%) and theophylline (0, 0.2, 0.5 mM). After incubation at 37°C for 3 days, cell growth was recorded.

### Spectrophotometric coaggregation assay

A previously published spectrophotometric coaggregation assay protocol was carried out (14). In brief, the bacterial cells were harvested, washed, and resuspended in phosphate-buffered saline (PBS). Equal numbers of cells were then diluted in the coaggregation buffer solution (150 mM NaCl, 1 mM Tris-HCl, 0.1 mM CaCl_2_, 0.1 mM MgCl_2_) to a final concentration of 2 x 109 cells/ml. After mixing the cell suspensions, samples were incubated for 10 minutes at room temperature. After incubation, the coaggregation reaction was centrifuged at low speed (100 g) for 1 minute to separate coaggregation cells from non-aggregated bacteria in suspension. The supernatant was carefully removed without disturbing the pellet, and its optical density was measured at 600 nm. The relative coaggregation of *F. nucleatum* (Fn) and A. oris (Ao) was determined by calculating the difference between the total turbidity of each partner strain and the coaggregation supernatant turbidity, divided by the total turbidity of each partner strain, using the formula of {[OD_600_(Fn) + OD_600_(Ao)]-OD_600_(Fn+Ao)}/[OD_600_(Fn) +OD_600_(Ao)].

### Electron microscopy

To prepare enough *lepB-depleted* cells for electron microscopy analysis, we grew an 8 ml overnight culture of Δ*lepB*pAN01 with 0.5% xylose and 1 mM theophylline. The overnight culture was centrifuged to obtain a cell pellet, which was washed three times with water to remove any residual inducers in the culture. Then the cell pellet was resuspended into 100 µl of 15% glycerol and kept at -80°C for overnight. The next day, the cell suspension was thawed and plated on the TSPC agar plate with and without inducers (Figure 5C). The plate was incubated at 37°C for 16 hours before harvesting cells. A loopful of cells for each plate was harvested and suspended in 0.1 M NaCl. A drop10 µl bacterial suspension drop was placed onto carbon-coated nickel grids and stained with 0.1% uranyl acetate. Samples were washed with water before imaging with a JEOL JEM1400 electron microscope. For analysis by scanning electron microscopy, fusobacterial cells were immobilized on a silicon wafer and fixed with 2.5% glutaraldehyde in PBS buffer (pH 7.3) for 1 hour at room temperature. After fixation, the samples were washed thoroughly with PBS buffer (pH 7.3). For dehydration, the samples were sequentially put in 20%, 40%, 60%, 90%, and 100% ethanol solution in water for respectively 15min. And then, the samples were put in a critical point dryer (Autosamdri-815 from Tousimis Research Corporation) in ethanol for 1.5h. After critical point drying, samples were coated with a thin iridium film of 8 nm with a magnetron sputtering Coater (208HR High-Resolution Sputter Coater, Ted Pella Inc imaged using a Nova NanoSEM 230 (FEI). The scanning work distance was 5 mm, and the accelerating high voltage was 5 kV.

## AUTHOR CONTRIBUTIONS

C.W. conceived and designed all experiments. B.G., P.Z., A.N., J.G and C.W. performed all experiments. B.G., P.Z., and C. W. analyzed data. C.W. wrote the manuscript with contribution and approval from all authors.

## DATA AVAILABILITY STATEMENT

Materials are available upon reasonable request with a material transfer agreement with UTHealth for bacterial strains or plasmids. Plasmids pBCG06 (190721) and pAN01 (190725) have been deposited with Addgene. Plasmids information can be accessed on Benchling links at https://benchling.com/s/seq-uREFKFed8I5A8RJtZDqh?m=slm-OgXchIQpBsKzdrXUQeA2(pBCG06); https://benchling.com/s/seq-QXreL05ooQTSsiJ10Ryp?m=slm-BYZe9zJsHuyPfj9ClNPY(pBCG06a); https://benchling.com/s/seqkPLE0WBATzQQMC1jEoIm?m=slm-XJyBJ3fBHNFZLN7NzzUp (pBCG07); https://benchling.com/s/seq-F6AxBucdWbSmgmNe3KPq?m=slm-NahcnWEgUhXPpcYlkTMw (pBCG08); https://benchling.com/s/seq-CniLnNtyGecEkv8efJVM?m=slm-2BdAC4H49hsPETNZU8sf (pAN01).

## ACKNOWLEDGMENTS

This work was supported by a National Institute of Dental and Craniofacial Research (NIDCR) grant (DE030895) to C.W. The plasmid pXCH was kindly provided by Dr. Hirofumi Nariya from Jumonji University in Japan. We thank Dr. Janina P. Lewis (Virginia Commonwealth University) for sharing the pG106 vector and Dr. Justin Merritt (Oregon Health & Science University) for providing an *S. mutant* strain JK-G-7 containing luciola red luciferase gene.

